# The influence of substrate concentration on the culturability of heterotrophic soil microbes isolated by high-throughput dilution-to-extinction cultivation

**DOI:** 10.1101/726661

**Authors:** Ryan P. Bartelme, Joy M. Custer, Christopher L. Dupont, Josh L. Espinoza, Manolito Torralba, Banafshe Khalili, Paul Carini

## Abstract

The vast majority of microbes inhabiting oligotrophic shallow subsurface soil environments have not been isolated or studied under controlled laboratory conditions. In part, the challenges associated with isolating shallow subsurface microbes may persist because microbes in deeper soils are adapted to low nutrient availability or quality. Here we use high-throughput dilution-to-extinction culturing to isolate shallow subsurface microbes from a conifer forest in Arizona, USA. We hypothesized that the concentration of heterotrophic substrates in microbiological growth medium would affect which microbial taxa were culturable from these soils. To test this, we diluted extracted cells into one of two custom-designed defined growth media that differed only by a 100-fold difference in the concentration of amino acids and organic carbon. Across both media, we isolated a total of 133 pure cultures, all of which were classified as Actinobacteria or Alphaproteobacteria. The substrate availability dictated which actinobacterial phylotypes were culturable but had no significant effect on the culturability of Alphaproteobacteria. We isolated cultures that were representative of the most abundant phylotype in the soil microbial community (*Bradyrhizobium* spp.) and representatives of five of the top 10 most abundant *Actinobacteria* phylotypes, including *Nocardioides* spp., *Mycobacterium* spp., and several other phylogenetically-divergent lineages. Flow cytometry of nucleic acid-stained cells showed that cultures isolated on low-substrate medium had significantly lower nucleic-acid fluorescence than those isolated on high-substrate medium. These results show that dilution-to-extinction is an effective method to isolate abundant soil microbes and the concentration of substrates in culture medium influences the culturability of specific microbial lineages.

**Importance:** Isolating environmental microbes and studying their physiology under controlled conditions is an essential aspect of understanding their ecology. Subsurface ecosystems are typically nutrient-poor environments that harbor diverse microbial communities—the majority of which are thus far uncultured. In this study, we use modified high-throughput cultivation methods to isolate subsurface soil microbes. We show that a component of whether a microbe is culturable from subsurface soils is the concentration of growth substrates in the culture medium. Our results offer new insight into technical approaches and growth medium design that can be used to access the uncultured diversity of soil microbes.

## Introduction

Soil microbial communities are tremendously diverse and mediate crucial aspects of plant fertility, biogeochemistry, pollutant mitigation, and carbon sequestration (1–4). While the diversity and community composition of surface soils have been relatively well described, we know far less about the microbes inhabiting deeper soils (defined here as >10 cm below the surface), despite their key role in soil formation and mineralization of key plant nutrients. In contrast to surface soils that are typically rich in plant-derived compounds, subsurface soils are often characterized by lower amounts of mineralizable nitrogen, phosphorus and organic carbon—much of which has a long residence time and is relatively recalcitrant to microbial degradation (5–9). The temperature and soil moisture of subsurface soils are also less variable than shallower soils that are exposed to seasonal changes in temperature and precipitation (10). These relatively stable and low-nutrient conditions found at depth constrain both the amount of microbial biomass present in the subsurface and the structure of these microbial communities (11–14). Many of the microbial taxa that are abundant in these subsurface environments are underrepresented in microbial culture and genome databases (11). Thus, there are large knowledge gaps in our understanding of the biology of a major fraction of subsurface soil microbes.

Because subsurface soils are low-nutrient habitats, part of the challenge associated with culturing and studying the microbes that live belowground may be that they require low nutrient concentrations in order to be isolated or propagated in the laboratory (15). These microbes— often referred to as ‘oligotrophs’—are capable of growing in conditions where the supply or quality of nutrition is poor. Although oligotrophs dominate most free-living microbial ecosystems (16), the concept of oligotrophy itself is enigmatic. There is no coherent definition of what constitutes oligotrophic metabolism aside from their ability to grow at ‘low’ nutrient concentrations—a definition that itself is arbitrary (17, 18). Kuznetsov *et al*. (19), identified three groups of cultivatable oligotrophs: 1) microbes that can be isolated on nutrient-poor medium, but cannot be subsequently propagated; 2) microbes that can be isolated on nutrient-poor medium but can be subsequently propagated on nutrient-rich medium; and 3) microbes that require special nutrient-poor medium for both isolation and propagation. Although the molecular and genetic mechanisms that distinguish these three categories are poorly understood, several traits of oligotrophs have emerged from the study of microbes that numerically dominate oligotrophic ecosystems. For example, oligotrophs are typically small, slowly growing cells (20–23). The genome sizes of numerous lineages of microbes that dominate oligotrophic marine ecosystems tend to be highly reduced—an indication that microbial oligotrophy may be tied to reduction of genome size (24–26). These ‘streamlined’ genomes often code for fewer copies of the rRNA gene operon and transcriptional regulator genes than microbes with larger genomes, suggesting oligotrophs lack the ability to sense and rapidly respond to varying environmental conditions (12, 16). Instead, genomic inventories of marine oligotrophs suggest a reliance on broad-specificity, high-affinity transporters that are constitutively expressed (22, 26, 28, 29).

While the activities of abundant and ubiquitous microbes that inhabit oligotrophic marine environments have been extensively investigated in recent years (24, 30, 31), far fewer studies have focused on the activities of microbes that dominate oligotrophic soil environments. Several soil studies used low-throughput techniques to show that reduced-nutrient solid media facilitated the isolation important soil microbes that were previously uncultured (32–35). While several agar-based high-throughput approaches have been used to isolate diverse microbes (36, 37), these approaches may not be appropriate to isolate microbes that thrive at micromolar amounts of growth substrate and do not form detectable colonies on solid media. Here, we adapt existing high-throughput dilution-to extinction protocols, originally developed for isolating abundant aquatic oligotrophic bacteria, to facilitate the isolation of soil microbes. We hypothesized that the concentration of heterotrophic growth substrates in a growth medium would constrain which taxa were able to be isolated on a custom-designed defined artificial medium. We tested this by extracting cells from oligotrophic subsurface soils using Nycodenz buoyant density centrifugation (38), and inoculating high-throughput dilution-to-extinction experiments in two defined media that contained a 100-fold difference in the amounts of heterotrophic growth substrates. We isolated several bacteria that were representative of abundant phylotypes found in the original soil microbial community and two lineages representative of uncultured groups of microbes. In these experiments, the substrate concentration significantly influenced which actinobacterial genera were culturable in the laboratory but no effect on which alphaproteobacterial lineages were culturable. Moreover, we show that cells isolated on low nutrient medium had significantly lower SYBR green I nucleic acid fluorescence, suggesting microbes isolated on low nutrient medium may contain reduced nucleic acid content relative to those isolated on higher nutrient medium.

## Results

We collected shallow subsurface soil (55 cm) from the Oracle Ridge field site in a mid-elevation conifer forest that is part of the Santa Catalina Mountains Critical Zone Observatory in Arizona, USA. These soils contained very low amounts of total organic carbon (0.095%) and N-NO^3^ (0.3 ppm) indicating they were highly oligotrophic (Supplementary Fig. 1). We adapted existing high-throughput dilution-to-extinction approaches designed for aquatic microbes (39, 40) to culture soil microbes from these samples (Fig. 1). The primary modification to existing protocols was to add a buoyant density centrifugation cell-separation step to detach inoculum cells from mineral soils prior to diluting cells into growth medium. To do this, we vortexed soil in a cell extraction buffer containing a nonionic surfactant and a dispersing agent. We layered this soil-buffer slurry over 80% Nycodenz and centrifuged it. During centrifugation, the mineral components of soil migrated through the Nycodenz, while cells ‘floated’ on the surface of the Nycodenz. We extracted cells located at the Nycodenz interface, stained them with SYBR Green I, and counted them on a flow cytometer. This extraction yielded 1.28 × 10^5^ cells ml^-1^ from 0.5 g wet soil. We diluted the extracted cells to an average of 5 cells well^-1^ in deep-well polytetrafluoroethylene 96-well plates containing a custom-designed and defined growth medium that we named ‘Artificial Subterranean Medium’ (ASM), with low or high concentrations of heterotrophic growth substrates (ASM-low and ASM-high, respectively) (Fig. 1). The ASM-low and ASM-high media contained identical inorganic mineral and vitamin amendments but a 100-fold difference in the concentration of organic carbon and amino acids (Supplementary Table 1). We designed these media to facilitate the growth of diverse chemoheterotrophic microbes by including an array of simple carbon compounds, polymeric carbon substrates, and individual amino acids (Supplementary Table 1). We prepared triplicate 96-well plates for each growth medium formulation. These dilution-to-extinction experiments were screened for growth with flow cytometry after 4 weeks of incubation and again after 11 weeks of incubation (Fig. 1). Wells displaying growth (defined as those wells displaying 1.0 × 10^4^ cells ml^-1^) were sub-cultured into larger volumes and subsequently cryo-preserved and identified by 16S rRNA gene sequencing (Fig. 1).

**Figure 1:**
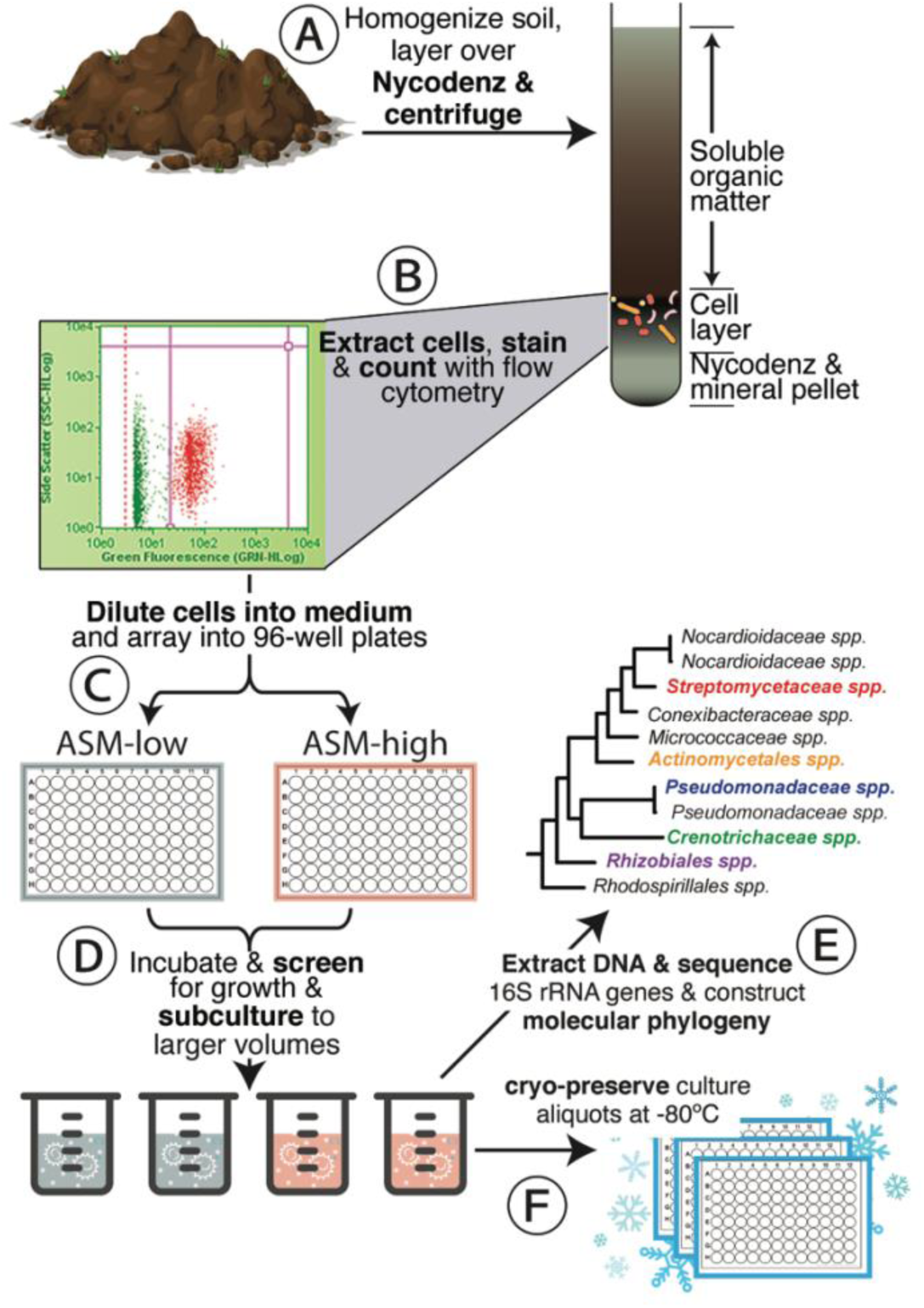
Dilution-to-extinction workflow. Soils were collected and brought to the lab where they were homogenized in cell extraction buffer, layered over a Nycodenz solution, and centrifuged (A). The cell layer was extracted from the Nycodenz solution and counted with flow cytometry (B). Counted cells were diluted into growth medium in 96-well microtiter plates to an average density of 5 cells well^-1^ (C). After incubation, the 96-well microtiter plates were screened for growth with flow cytometry, and wells displaying growth were subcultured into larger volumes (D). After incubating the subcultures, flasks displaying growth were identified by 16S rRNA gene sequencing and molecular phylogeny (E). Aliquots of these identified subcultures were cryopreserved at -80°C (F).

Across both medium types, a total of 214 wells (119 for ASM-low and 95 for ASM-high) displayed growth after 11 weeks of incubation. We successfully propagated 182 (85%) of the cultures from microtiter plates to polycarbonate flasks containing fresh medium. Of the cultures that successfully propagated, we confirmed that 73% (133 cultures) were pure cultures by amplifying and sequencing full-length 16S rRNA gene sequences from genomic DNA extractions. The remaining 49 cultures were mixed (Foreword and Reverse 16S rRNA sequence reads did not assemble due to base ambiguities) or, in rare instances, did not amplify under several amplification conditions. We defined microbial culturability using Button’s definition of microbial ‘viability’ as determined in dilution-to-extinction experiments (41). Here, ‘culturability’ is defined as the ratio of cells that grew into detectable cultures to the total number of cells initially diluted into a cultivation chamber (41). By the end of the experiment, we approached ∼20% culturability across both medium formulations (Fig. 2). In general, microbial culturability was higher for ASM-low than for ASM-high, but this effect was only significant after 4 weeks of incubation (Fig. 2; Wilcoxon rank-sum test *P* ≤ 0.05 at 4 weeks).

**Figure 2:**
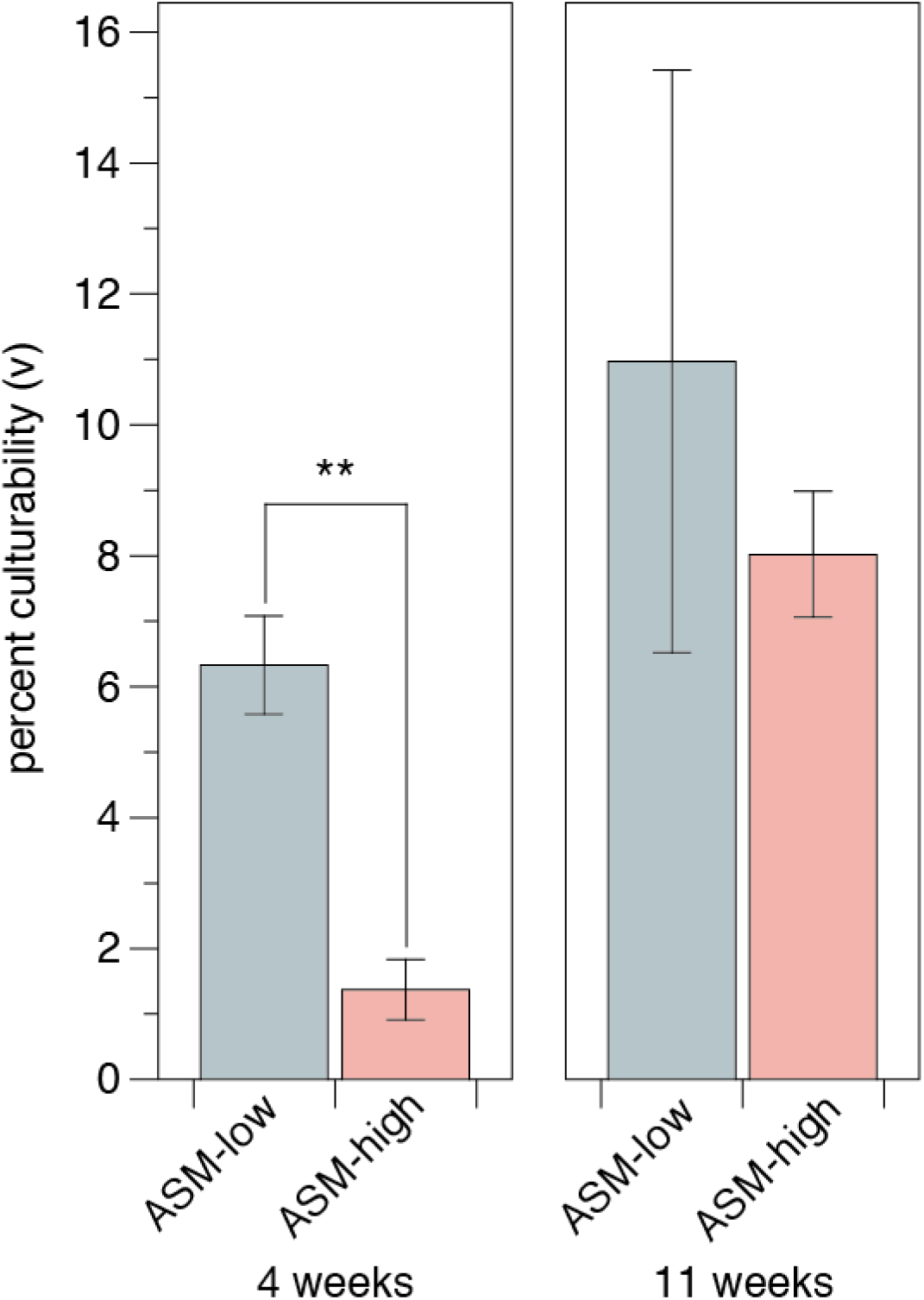
Microbial culturability (v) was greater on ASM-low than on ASM-high. Bar heights are the mean percent culturability ± standard deviation in 96 well microtiter plates (n=3) as calculated from the initial cell inoculum and the proportion of wells positive for growth (41). Double asterisks indicate Wilcoxon rank sum test P values ≤ 0.05.

We assigned taxonomy to each 16S rRNA gene sequence using the SILVA database. All pure cultures isolated on ASM-low and ASM-high belonged to one of two bacterial phyla: Actinobacteria (110 cultures; 83% of the pure cultures) or Proteobacteria (23 cultures, all Alphaproteobacteria; 17% of the pure cultures) (Fig. 3). Across all experiments, the genera assigned to bacteria isolated on ASM-low were significantly distinct from those isolated on ASM-high (Kruskal-Wallis rank sum, χ^2^=19.05, *P*=1.28 × 10^-5^). However, these differences were largely driven by significant differences in culturability across medium types for the Actinobacteria, but not for the Alphaproteobacteria (Dunn test, *P*≤0.000 for Actinobacteria and *P*=0.128 for Alphaproteobacteria).

**Figure 3:**
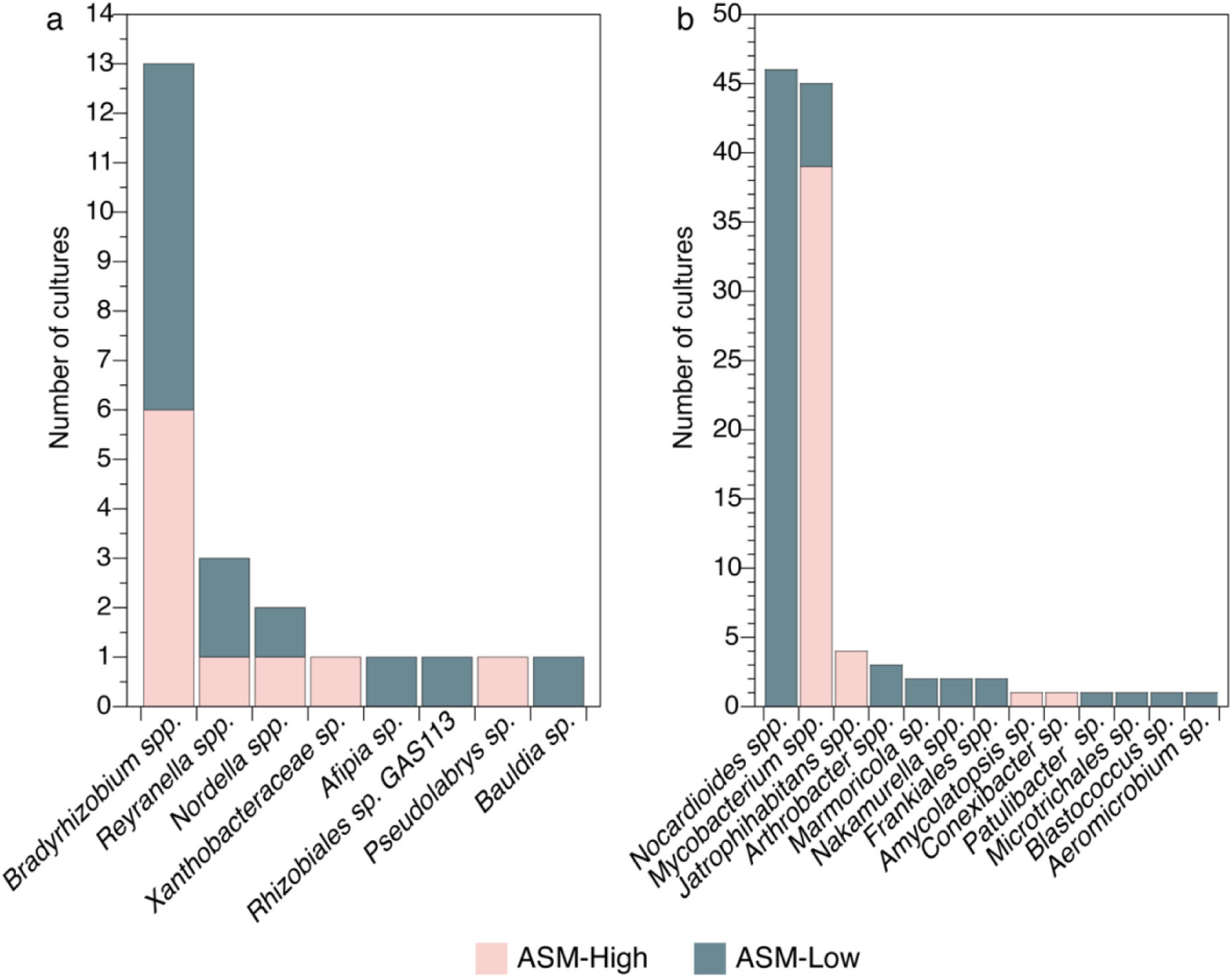
ASM-low and ASM-high cultured distinct Alphaproteobacteria (a) and Actinobacteria (b). Bar heights are the number of cultures obtained for each taxon and are colored by the medium type on which they were isolated. The genera assigned to bacteria isolated on ASM-low were distinct from those isolated on ASM-high (Kruskal-Wallis rank sum, χ^2^=19.05, *P*=1.28 × 10^-5^). These differences were driven by differences in culturability across medium types for actinobacterial genera, but not for alphaproteobacterial genera (Dunn test, P≤0.000 for Actinobacteria and P=0.128 for Alphaproteobacteria).

Just over half of the Alphaproteobacteria (57%) were isolated on ASM-low medium and the remaining 43% on ASM-high (Fig. 3 and Supplementary Fig. 2). Cultures that classified as *Bradyrhizobium* spp. were the most frequent alphaproteobacterial isolates (13 isolates), seven of which were isolated on ASM-low medium. Cultures classified as *Reyranella* spp. and *Nordella* spp. were also isolated on both ASM-high and ASM-low medium. Of the remaining five Proteobacteria cultures, three were isolated on ASM-low (*Afipia* (1 culture), *Rhizobiales* (1 culture), *Bauldia* (1 culture)), and two were isolated on ASM-high (*Pseudolabrys* (1 culture) and a *Xanthobacteraceae* sp. (1 culture)). The actinobacterial cultures belonged to three classes: Actinobacteria (107 cultures), Thermoleophilia (2 cultures), and Acidimicrobiia (1 culture). Of these Actinobacteria, 65 (59%) were isolated on ASM-low, and 45 (41%) on ASM-high. The cultures were numerically dominated by two genera that were differentially isolated on ASM-low and ASM-high: *Nocardioides* and *Mycobacterium*. *Nocardioides* spp. (46 cultures) were exclusively isolated on ASM-low medium (Fig. 3 and Supplementary Fig. 3). Other cultures that were isolated on ASM-low included those classified as *Arthrobacter* (3 cultures), *Marmoricola* (2 cultures), *Nakamurella* (2 cultures), *Aeromicrobium* (1 culture), *Blastococcus* (1 culture), and *Patulibacter* (1 culture) (Fig. 3 and Supplementary Fig 3). While the majority of cultures classified as *Mycobacterium* sp. were isolated on ASM-high (38 cultures), we isolated seven mycobacterial cultures on ASM-low medium—five of which form a phylogenetically-distinct cluster from those isolated on ASM-high (Fig. 3 and Supplementary Fig. 3). Other actinobacterial cultures isolated on ASM-high included *Jatrophihabitans* (4 cultures), *Conexibacter* (1 culture), and *Amycolatopsis* (1 culture).

Interestingly, we isolated what are likely the first members of two novel actinobacterial lineages on ASM-low. The first such culture—*Microtrichales* sp. str. AZCC_0197—belongs to the *Microtrichales* order of the Acidimicrobiia class. The best 16S rRNA gene sequence match to an existing isolate is 93.4% identity to *Aquihabitans daechunggensis* str. G128. However, strain AZCC_0197 more closely matched numerous 16S rRNA gene sequences from environmental clones of uncultured Acidimicrobiia. The second lineage—*Frankiales* sp. strains AZCC_0102 and AZCC_0072—classified as members of the *Frankiales* order of the Actinobacteria class with best matches of <97% nucleotide identity to existing *Frankiales* isolates (42).

Several of the microbes we isolated were representative of abundant members of the subsurface soil microbial community at the Oracle Ridge site. We matched the 16S rRNA gene sequences from our cultures to the phylotypes derived from the 55 cm Oracle Ridge soil sample. The 16S rRNA gene sequences from our cultures matched 13 phylotypes (at 97% identity; Fig. 4) that account for 11.0 ± 1.6% (mean ± SD, n=3) of the total amplifiable microbial community. For example, the 16S rRNA gene sequences from our *Bradyrhizobium* isolates match a single *Bradyrhizobium* phylotype that was the most abundant phylotype at 55 cm (relative abundance of 5.7 ± 0.3% (mean ± SD, n=3); Fig. 4). Additionally, we isolated representatives of abundant Actinobacteria (Fig. 4) including: two *Mycobacterium* phylotypes (the 11^th^ and 17^th^ most abundant phylotypes overall); *Nocardioides* (the 13^th^ most abundant phylotype overall); and two *Arthrobacter* phylotypes (16^th^ and 1,271^st^ most abundant phylotypes overall). The other Actinobacteria cultured in these experiments represent rarer phylotypes in bulk soils. The 16S rRNA gene sequences from several of our pure cultures did not match any of the phylotypes derived from these soils at ≥97% identity, including *Nakamurella* (2 cultures)*, Nocardioides* (5 cultures), *Mycobacterium* (1 culture), *Jatrophihabitans* (1 culture), *Patulibacter* (1 culture), *Conexibacter* (1 culture), *Rhizobiales* sp. (1 culture), *Reyranella* (1 culture), and *Microtrichales* sp. str. AZCC_0197.

**Figure 4:**
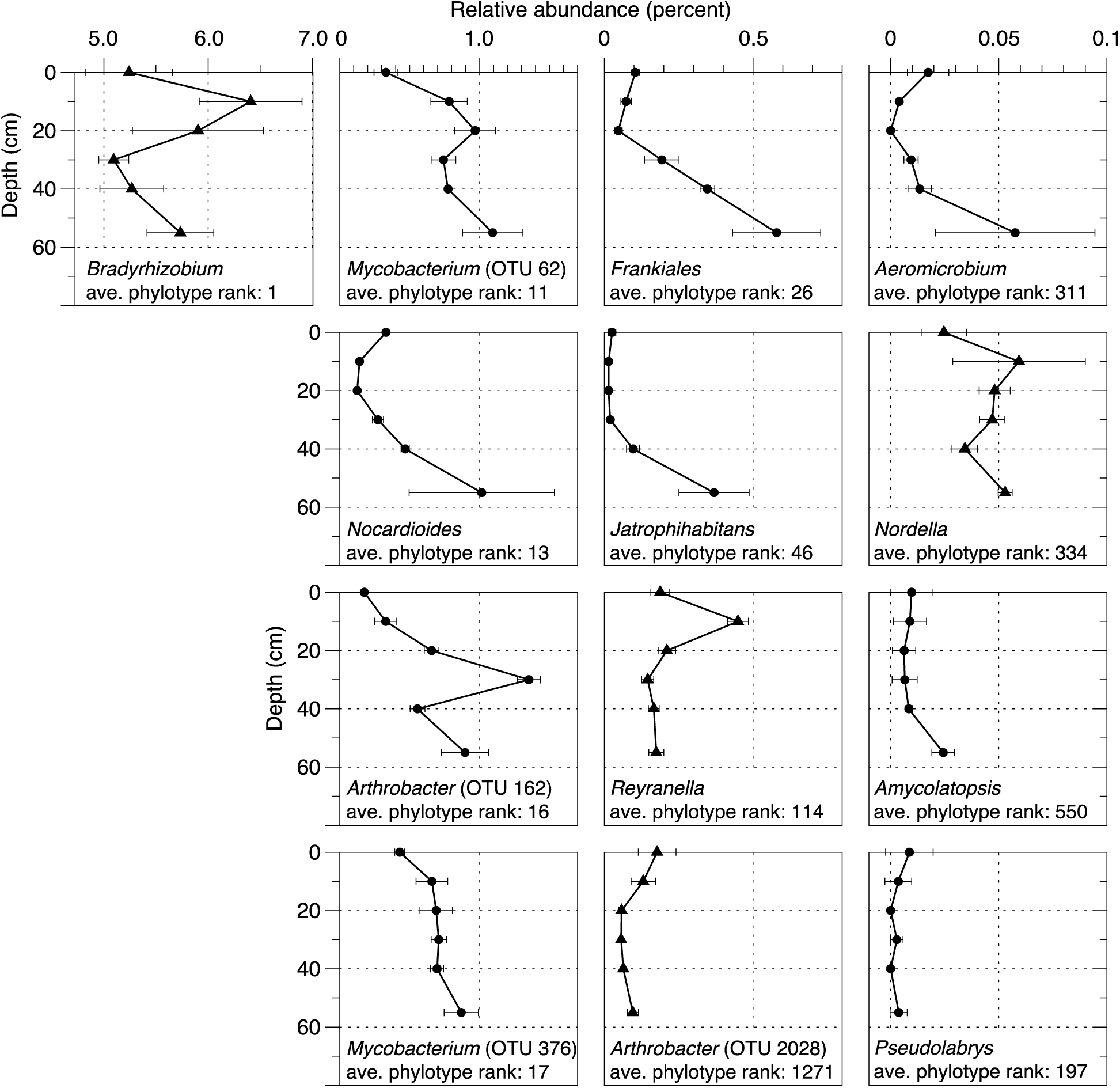
The cultures isolated in this study were representative of several abundant soil lineages that show dynamic depth distributions in Oracle Ridge soils. Points are the mean relative abundances ± standard deviation (n=3) of 16S rRNA gene sequence phylotypes that matched the 16S rRNA gene sequences obtained from cultured isolates at ≥97% identity. Error bars that are not visible are located behind the symbol. Assigned genera names and the average (n=3) relative rank of each phylotype at 55 cm are shown. Cultures classified at the genera level as *Mycobacterium* and *Arthrobacter* cultures matched more than one phylotype in the cultivation-independent surveys. The best-matching OTU number is designated in parentheses. Triangles are Alphaproteobacteria. Circles are Actinobacteria.

In many environments, relative nucleic acid content can be estimated with flow cytometry analysis of cells stained with nucleic acid staining dyes (43–45). Given that many microbes inhabiting low-nutrient environments exhibit reduced genome sizes, we sought to determine whether the nucleic acid fluorescence measured for our cultures partitioned by the growth medium they were isolated on. For each culture, we identified the closest match to an available genome sequence and found that the genome length of these ‘best hit’ matches was significantly correlated with the average fluorescence of SYBR Green I-stained cells (Spearman’s rho=0.3 *P*=0.0012), indicating that the nucleic acid fluorescence we quantified by flow cytometry was indeed indicative of genome size differences. We found that cultures isolated on ASM-low exhibited significantly lower mean nucleic acid fluorescence than those isolated on ASM-high (Fig. 5; Kruskal-Wallis Rank sum χ^2^=24.8 *P*=6.27 × 10^-7^). The overall mean fluorescence was not significantly different across the phyla assigned to each isolate (Kruskal-Wallis Rank sum χ^2^=0.210, *P*=0.647), but was significant across individual genera assignments (Kruskal-Wallis Rank sum χ^2^=62.4 *P*=2.98 ×10^-6^). Moreover, the mean nucleic acid fluorescence values within a given genera were similar (Supplementary Fig. 4). For example, *Mycobacterium* isolates had relatively high nucleic acid fluorescence, regardless of which medium they were isolated on (Supplementary Fig. 4). In contrast, *Nocardioides* (ASM-low) and *Jatrophihabitans* (ASM-high) displayed relatively low nucleic acid fluorescence. Interestingly, when cultured on ASM-high, we observed a clear nucleic acid fluorescence dichotomy across the *Bradyrhizobium* isolates (Supplementary Fig. 4).

**Figure 5:**
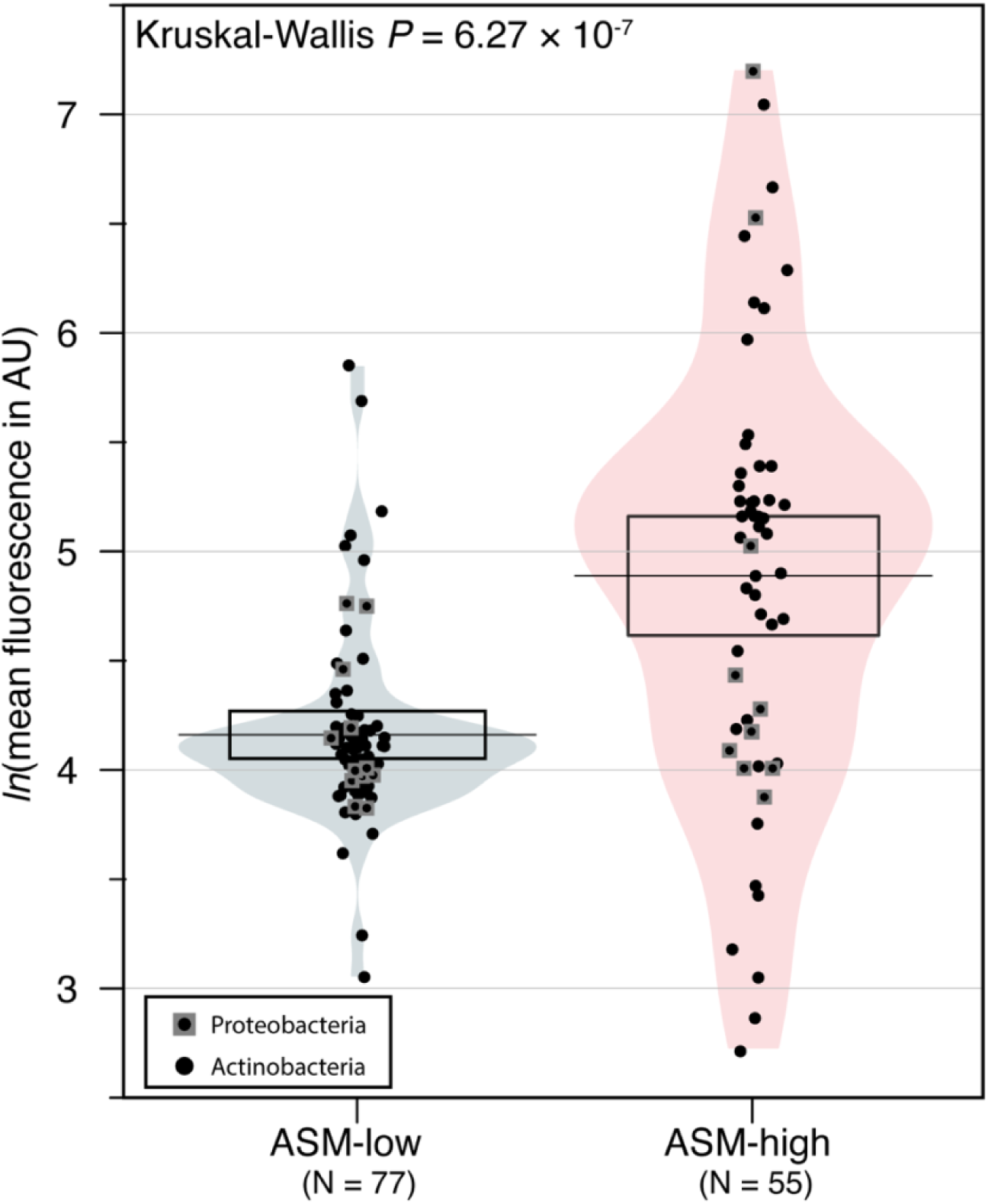
The mean nucleic acid fluorescence of taxa isolated on ASM-low was significantly lower than for those microbes isolated on ASM-high. Points are the mean natural logarithm (ln) of the quantified nucleic acid fluorescence (in arbitrary units [AU]) of fixed and SYBR green I-stained stationary phase cultures. The mean fluorescence value was obtained from manually-gated histogram plots of fluorescence within the Guava EasyCyte software. Only those cultures that were defined as pure cultures are plotted. The horizontal line in each plot is the mean fluorescence value and the box surrounding the mean is a 95% confidence interval. Shading illustrates the relative distribution of fluorescence values within each medium type.

## Discussion

We designed a proof-of-concept workflow to determine the feasibility of high-throughput dilution-to-extinction cultivation for the isolation of soil microbes. The method was based on a workflow for isolating microbes from oligotrophic marine environments (39). However, unlike aquatic samples, microbial cells in soils are heterogeneously dispersed within, or attached to, a complex matrix comprised of non-cellular organic matter and minerals. The complexity of this soil matrix complicates accurate enumeration of viable cells because mineral and organic matter can interfere with flow cytometry. To circumvent these issues, we separated cells by gently shaking soils in a cell extraction buffer containing a dispersing agent and a nonionic surfactant. Cells were separated from this slurry by buoyant density centrifugation (Fig. 1). This procedure allowed cells to be floated on top of a dense solution of Nycodenz while allowing minerals to migrate through the Nycodenz solution (46).

We estimated the expected culturability and calculated the actual culturability using the statistical framework of dilution culture growth outcomes described by Button, *et al.* (41). The expected number of pure cultures (*û*) was estimated across all experiments using the formula *û* =-*n*(1-*p*)×ln(1-*p*). Where *p* is the proportion of wells displaying growth (214 growth chambers displaying growth ÷ 576 chambers inoculated = 0.37) and *n* is the number of inoculated chambers (576 chambers in total). Based on this equation, we expected *û* = 168 pure cultures across all experiments. The number of pure cultures we obtained (133 cultures) was within 21% of this value. However, this result is conservative because it does not account for the cultures that were initially scored as positive for growth but did not successfully subculture. Some of these cultures may be oligotrophic taxa that were initially cultivatable but failed to successfully propagate, as described by Kuznetsov et al. (19). Alternatively, cultures that failed to propagate from microtiter plates to larger volumes might have been false positives, where flow cytometer instrument noise or well-to-well carryover was mistaken for a low-density culture. The mean culturability we observed for a given experiment (1.4%-11%; Fig. 2) was comparable to dilution-to-extinction cultivation studies of marine microbes, which report 0.5%-14.3% culturability (40). Similar to previous observations for soil microbes (34), we observed increased culturability with longer incubation times (Fig. 2). We speculate that the culturability was higher on ASM-low than on ASM-high because the cells were extracted from an extremely oligotrophic soil sample (Supplementary Fig. 1) and perhaps the results would have been different if the original inoculum originated from more productive soils.

The concentration of heterotrophic growth substrates in the isolation medium significantly influenced our ability to isolate certain actinobacterial lineages (Fig. 3). In particular, *Nocardioides* were exclusively isolated on ASM-low and most *Mycobacterium* isolates were isolated on ASM-high. We designed our media to include a defined but diverse range of carbon substrates that have been successfully used to isolate chemoheterotrophic microbes from soil or oligotrophic taxa from other environments (34, 50). Carbon type and availability are crucial for heterotrophic soil microbes because carbon acts as both a source of electrons for respiration and carbon for biomass. To accommodate this requirement, many common microbial growth medium formulations for heterotrophic microbes supply diverse growth substrates (yeast extract or casein digests, for example), usually at concentrations much higher than are normally available *in situ*. Two key assumptions made with these common media formulations are that: 1) microbes will use only the relevant constituents and any remaining compounds will have minor or no effect on microbial growth; and 2) microbes grow optimally in the laboratory when nutrient availability is much greater than their half saturation constant (47). While many commonly studied microbes have the capacity to grow on complex, high-nutrient formulations, environmental nucleic acid data informs us that the vast majority of Earth’s microbes remain uncultured (48, 49). Our results indicate that the concentration of nutrients in a growth medium may be as important as the constituents of the growth medium for cultivation of uncultured environmental microbes.

Numerous studies have demonstrated that dilute growth media is superior to substrate rich growth media for the isolation of novel soil microbes (34, 35, 51, 52). However, the physiological explanation as to why low-nutrient media facilitates the growth of diverse microbes, or high nutrient concentrations inhibit the growth of some taxa, remain unclear. One possible explanation for these concentration-dependent effects may be that growth medium formulations applied at high concentrations contain high amounts of inhibitory substances— substances that are reduced to non-inhibitory levels in dilute medium formulations. For example, a key amino acid transporter in *Chlamydia trachomatis* can be blocked by non-essential amino acids, preventing the transport of required amino acids, resulting in growth inhibition (53). A similar phenomenon was demonstrated in the extreme marine oligotroph ‘*Candidatus* Pelagibacter ubique’, where alanine was conditionally required for cell division but abolished growth at higher concentrations (54). Furthermore, reactive oxygen species can be produced during the autoclaving of nutrient-rich medium that either directly inhibit growth or combine with organics in the medium to form inhibitory compounds (55, 56). Finally, growth inhibition may be the result of misbalanced regulation of growth or accumulation of nutrient storage structures (poly-β-hydroxybutyrate, for example) ultimately leading to cell lysis (57). A better understanding of the mechanism(s) that enable growth on low nutrient medium—or prevent growth on high-nutrient medium—may help us design better strategies for isolating uncultivated lineages. Critically, the collection of cultures we describe here, that were isolated on medium with identical constituents applied at different concentrations, is a first step towards an experimental method capable of addressing these questions.

While several of the taxa we isolated were abundant microbial members of the shallow subsurface microbial community (Fig. 4), other isolates were rare or not identified in the cultivation-independent soil microbial community. The cultivation of additional microbial phylotypes that were not observed in molecular analyses of the same samples has been observed (58, 59). The dilution-to-extinction approach we used here favors the cultivation of abundant microbes in a given sample (41), such that the isolation of rarer taxa or taxa that were not observed in the original sample was unexpected. There are several possible explanations for this observation. First, the buoyant density separation protocol that we used to separate cells may have introduced biases. Nycodenz cell separation approaches do not recover all microbial cells from soil and the separable fraction can be compositionally distinct from the non-separated soils (61). These effects could potentially skew the proportions of microbes that were diluted into microtiter plate wells. Alternatively, the absence of a particular taxon in a soil microbiome analysis may be the result of insufficient sequencing depth (58). Moreover, the ‘universal’ primers used in the soil microbiome analysis (60) may not have primed DNA from some of the divergent lineages we cultured as efficiently as other phylotypes in the soil microbiome, resulting in either underrepresentation of these phylotypes in the original community or no amplification at all. We do not have sufficient evidence indicating which of these scenarios may explain our ability to culture cells that were not apparent in the microbiome analysis. Finally, as is true in any microbial cultivation experiment, there were many taxa that we did not isolate. In particular, these soils contained high relative abundances of Verrucomicrobia related to ‘*Candidatus* Udaeobacter copiosus’ (62), and Acidobacteria (Subgroup 6), which belong to highly sought-after lineages of uncultured microbes (63). The reasons for not isolating these (and other) lineages are numerous but may be the result of inappropriate medium composition (64, 65), toxic compounds in the cell separation constituents, long doubling times (> ∼6 days), or dormancy (reviewed in (66)).

We provide evidence that microbes cultured from oligotrophic soils on low-nutrient medium may have reduced nucleic acid content relative to microbes isolated on richer medium (Fig. 5). Depending on the taxa in question, and their effective population size, microbial genome reduction can be driven by either genetic drift or ‘streamlining’ selection (reviewed in (67)). Genome streamlining is strongly linked with microbial oligotrophy in free-living aquatic microbes as a mechanism to reduce the overhead cost of replication in periodically nutrient-limited environments (reviewed in (21)). However, direct evidence for genome streamlining in terrestrial microbes has been elusive. For example, metagenome-assembled genomes of abundant and ubiquitous uncultured Verrucomicrobia suggest that some lineages may contain reduced genome sizes (62). A more recent study showed that fire-affected warm soils selected for groups of microbes with significantly smaller genomes than cooler soils (68). Yet, there are few definitive ways to identify the growth preferences of taxa with reduced genomes short of culturing them and studying their growth dynamics under controlled conditions. The appearance of reduced nucleic acid content in cultures isolated on ASM-low may be an indication that genome reduction may be a successful life strategy for soil oligotrophs. Alternative explanations for the apparent differences in nucleic acid content in microbes cultured in ASM-low may be that: 1) the ploidy of stationary-phase cells grown in ASM-low may be lower than those isolated in ASM-high; 2) unknown cellular constituents of cells grown in ASM-low may quench SYBR Green I fluorescence in the assay conditions we used; or 3) cells isolated in ASM-high may form small microaggregates that are not completely dispersed prior to flow cytometry.

The development of cultivation techniques emphasizing the high-throughput and sensitive detection of microbial growth on low-nutrient medium revolutionized the field of aquatic microbial ecology by culturing microbes that were previously ‘unculturable’ using standard techniques (39, 40, 50, 69, 70). Here we show that similar cultivation principles facilitate the cultivation of abundant soil microbes. We demonstrate that, in addition to scrutinizing the nutritional composition of a given growth medium, the concentration of growth substrates in the growth medium must also be considered. Although we do not yet understand the mechanism of substrate-induced growth inhibition, there is evidence that this phenomenon is widespread and may impede laboratory cultivation efforts. Future studies to deduce the molecular mechanisms of substrate-induced growth inhibition will likely lead to new cultivation approaches that will allow us to isolate abundant free-living oligotrophic microbes.

## Materials and Methods

### Soil source and nutrient analysis

Fresh soil samples were collected from a soil pit on August 16, 2017 at the Oracle Ridge site in the Catalina Jemez Critical Zone Observatory (coordinates: 32.45 N, -110.74 W, elevation 2,103 m). The mean annual precipitation at Oracle Ridge is 87 cm year^-1^ and the mean annual surface temperature is 12°C (71). The soils were Typic Ustorthent (71). The dominant vegetation at the site is ponderosa pine (*Pinus ponderosa*), with sparse Douglas-fir (*Pseudotsuga menziesii*). We collected ∼300 g subsamples from 0-5 cm, 10 cm, 20 cm, 30 cm, 40 cm and 55 cm depths. Soils were kept cool with ice packs for <4 h while in transit to the laboratory. At the laboratory, the soils were sieved to 2 mm and kept at 4°C for 50 days at which point cells were separated from mineral soil. Standard soil chemical analyses were performed at the Colorado State University Soil Water and Plant Testing Laboratory using their standard protocols. We analyzed the microbial community composition at each depth (see *‘Soil microbial community analysis,’* below) and conducted cultivation experiments from the 55 cm soil sample.

### Soil microbial community analysis

We extracted DNA from 1.0 g subsamples (n=3) using MoBio PowerSoil DNA extraction kits using the manufacturer’s instructions. We amplified 16S rRNA gene fragments using 515F-Y (5’-TATGGTAATTGTGTGYCAGCMGCCGCGGTAA-3’) and 926R (5’-AGTCAGTCAGGGCCGYCAATTCMTTTRAGT-3’) (72). PCR products were purified using the QIAquick PCR purification kit (Qiagen, Germantown, MD) per manufacturer’s specification. Cleaned products were quantified using Tecan fluorometric methods (Tecan Group, Mannedorf, Switzerland), normalized, and pooled for Illumina MiSeq sequencing using custom sequencing primers and the MiSeq Reagent v2 500 cycle Kit (Roche, Branford, CT) following the manufacturer’s protocols. We identified phylotypes based on the generation of *de novo* operational taxonomic units (OTUs) from raw Illumina sequence reads using the UPARSE pipeline at a stringency of 97% identity (73). Paired-end reads were trimmed of adapter sequences, barcodes and primers prior to assembly. We discarded low-quality and singleton sequences and dereplicated the remaining sequences before calculating relative abundances. Chimera filtering of the sequences was completed during clustering while taxonomy was assigned to the OTUs with mothur (74) using version 123 of the SILVA 16S ribosomal RNA database (75) as the reference. We generated OTU and taxonomy assignment tables for subsequent analyses.

### Cell separation

Cells were separated from sieved soils using buoyant density centrifugation with Nycodenz modified from (38) to isolate viable cells. Briefly, we added 0.5 g wet soil to 44.8 ml of cell extraction buffer (137.5 mM NaCl, 26.78 mM tetrasodium pyrophosphate, and 0.27% (v/v) Tween 80). The soil-buffer slurry was vortexed for 30 seconds and shaken horizontally on a platform shaker for 2 hours at 4°C. We layered 15 ml aliquots of this soil-buffer slurry over 10.0 ml of 80% (w/v) Nycodenz solution in 50 mM tetrasodium pyrophosphate. We used 50 ml Nalgene Oak Ridge high-speed polycarbonate centrifuge tubes for buoyant density centrifugations. Tubes containing the soil-buffer solution with Nycodenz were centrifuged at 17,000 × *g* for 30 minutes at 16°C. We extracted 3 × 0.5 mL aliquots from the resulting buoyant density preparation at a location of ∼25 mm above the bottom of the tube (coincident with the approximate level of the top of the Nycodenz solution) to sterile microcentrifuge tubes containing 1.0 ml 137.5 mM NaCl. The microcentrifuge tubes containing Nycodenz/NaCl were vortexed and centrifuged for 20 min at 17,000 × *g*. The resulting cell pellets were resuspended in 137.5 mM NaCl, pooled and stored at 4°C.

### Medium design rationale

The ASM media were custom-designed to facilitate the growth of a broad range of soil chemoheterotrophic microbes (Supplementary Table 1). Both ASM-high and ASM-low were buffered with phosphate. To this, we added minerals at concentrations derived from an ‘artificial rainwater’ recipe (76), trace elements as described in trace element solution SL-10 with the addition of LaCl^3^, and vitamins as described elsewhere (54). We added heterotrophic growth substrates that included 21 amino acids and a diverse range of simple carbon substrates including 2-C substrates (glycerol, acetate), 3-C substrates (pyruvate), 4-C substrates (succinate, butyrate, isobutyrate), 5-C substrates (ribose, valerate), a 6-C substrate (glucose), an 8-C substrate (NAG) and a 10-C substrate (decanoic acid) (Supplementary Table 1). We also added several polymeric growth substrates including pectin, methylcellulose, alginate, starch, and xylan (Supplementary Table 1). We calculated the added carbon amount to be ∼200 mg C L^-1^ for ASM-high and ∼2 mg C L^-1^ for ASM-low.

### Dilution-to-extinction

An aliquot of cells extracted from the buoyant density separation were fixed with 1.75% (final v/v) formaldehyde and stained with SYBR Green I (final stain concentration was a 1:4,000 dilution of commercial stock) for 3.5 h at room temperature in the dark. Cells were enumerated using a Millipore Guava flow cytometer, as described elsewhere (54). We diluted cells into Artificial Subterranean Medium (ASM)-high or ASM-low nutrient medium (Supplementary Table 1) to a density of 5 cells mL^-1^ and aliquoted 1.0 ml of the dilute cell suspension into the wells of 2 ml polytetrafluoroethylene 96-well microtiter plates (Cowie Technology, New Castle, DE) so that on average each well contained 5 cells. Plates were covered with plastic lids that allow air circulation and incubated at 16°C in the dark in aerobic conditions. We screened the dilution-to-extinction plates for growth by fixing (1.75% formaldehyde) and staining (1:4,000 dilution of commercial SYBR Green I stock) aliquots for 18 h in the dark at room temperature and counting by flow cytometry (EMD-Millipore Guava EasyCyte), as described previously (54). We screened plates for growth at 4 and 11 weeks after inoculation. Positive cultures were defined as cultures that exceeded 1.0 × 10^4^ cells ml^-1^.

### Actual and theoretical culturability estimates

Culturability estimates were determined by the equation: *V* = -ln(1-*p*)/*X,* where *V* is the estimated culturability*, p* is the proportion of inoculated cultivation chambers that displayed measurable growth (number of chambers positive for growth ÷ total number of chambers inoculated), and *X* is the number of cells added to each cultivation chamber as estimated from dilutions (41). The number of pure cultures (*û*) was estimated as follows: *û = -n(1-p) × ln(1-p)*, where *n* is the number of inoculated growth chambers and *p* is the proportion of inoculated wells displaying growth (41).

### Culture transfer and storage

We subcultured positive growth chambers into 25 ml of the respective growth medium (ASM-high or ASM-low) in acid-washed, sterile polycarbonate flasks and incubated them at 16°C. At the time of transfer, we assigned cultures a unique Arizona Culture Collection (AZCC) number. Flasks were monitored for growth every other week for 2 months. Flasks displaying growth within two months were cryopreserved in 10% glycerol and stored at -80°C. If no growth appeared within two months, the cultures were discarded and the assigned AZCC number was retired.

### Mean fluorescence calculations

We calculated the mean fluorescence of each culture from the subcultures grown in 25 ml volumes at 12-15 weeks after inoculating. Culture aliquots were fixed and stained for 15-18 h as described above in ‘Dilution-to-extinction.’ We manually gated histograms of the intensity of SYBR Green I fluorescence (in arbitrary units) and extracted the mean fluorescence of the gated peak for each culture using the GuavaSoft software package. ‘Best hit’ genomes were determined by blasting the full length 16S rRNA gene sequence of our isolates against the NCBI Microbial Genomes database using web-blast (77). We extracted the total genome length from each best-hit genome.

### Culture identification

Cultures were identified by full length 16S rRNA gene sequencing. Briefly, we filtered 5-10 ml of cell biomass from 25 mL cultures on to 0.2 µm-pore size Supor filters and extracted DNA using a Qiagen PowerSoil DNA extraction kit following the manufacturer’s instructions. We amplified full-length 16S rRNA genes from the resulting DNA using the 27F-1492R primer set (27F: 5’-AGAGTTTGATCMTGGCTCAG-3’; 1492R: 5’-ACCTTGTTACGACTT-3’ (78)). The reaction mix consisted of Promega’s GoTaq HotStart 2x PCR master mix with final concentrations of 0.4 µM 27F and 0.4µM 1492R primers, and 1-11.5 µl of template DNA, in a total reaction volume of 25 µL. The thermocycling profile was 1 × 94°C for 10 min followed by 36 cycles of: 94°C for 45 s, 50°C for 90 s, 72°C for 90 s; and a single 72°C extension for 10 min. The resulting amplicons were cleaned and Sanger sequenced from both the 27F and 1492R primer by Eurofins Genomics (Louisville, KY, USA) using their standard techniques. Sequences were curated using 4Peaks (79) and Geneious Prime v2019.0.1 (80). Reads were trimmed and assembled using the moderate setting in Geneious. Forward and reverse Sanger PCR reads that failed to build a full length 16S rRNA gene with these metrics were considered “mixed” cultures and not analyzed further.

### Culture taxonomy and determination of taxonomic differences across growth medium formulations

High-quality full length 16S rRNA gene sequences from the cultures were used to assign taxonomy and reconstruct phylogenetic relationships. We assigned taxonomy to all assembled 16S rRNA gene sequences using the SILVA database SINA aligner v128 (81). A Shapiro-Wilk test of normality was conducted in base R (82) on the distribution of SILVA genera assignments from both media types. After concluding the data were non-parametric, we performed a Kruskal-Wallis test in R (Assigned Genus ∼ Media Type). We performed a *post hoc* analysis (Dunn test, in R) to determine whether culturability within a Phylum varied by growth medium type.

### Taxonomic selection for phylogenetic reconstruction

To reconstruct a phylogeny of full length 16S rRNA genes, our culture sequences were compared to NCBI’s Microbial Genomes and environmental sequence databases using web-blast (77). The top five hits for each sequence from each NCBI database were chosen based on the highest percent coverage and lowest e-value score and included in the reconstruction. *E. coli* K-12 was used as the outgroup of the Alphaproteobacterial phylogeny, and *Bacillus subtilis* was used as the outgroup for the Actinobacterial tree. These sequences aligned with MAFFT (83) with turn checking enabled. The alignment was then trimmed using TrimAl (84) with the ‘automated1’ setting to optimize sequence trimming for maximum likelihood phylogenetic analyses. We reconstructed phylogenetic relationships from this trimmed alignment in the CIPRES Gateway (85).

Maximum-Likelihood (ML) trees were constructed using IQ-TREE with 10,000 ultrafast bootstrap trees and Bayesian Information Criterion to select the best fit nucleic acid substitution model (86, 87). For Actinobacteria, we used the SYM+R10 model and for Alphaproteobacteria we used the GTR+F+I+G4 model. After an initial round of ML trees, sequence alignments were heuristically curated with IQ-TREE to eliminate sequences that appeared in the tree more than once. Finalized ML trees were then imported into the ARB environment (88), where any duplicate sequences from our AZCC cultures were added to the ML trees through ARB’s quick add parsimony function. Final trees were visualized with FigTree (89).

### Environmental contextualization of AZCC isolates

We matched the AZCC isolate full length 16S rRNA gene sequences against a database of the clustered OTUs from the shallow soil depth profile samples (see *‘soil microbial community analysis,’* above) using the usearch_global command (90) at a stringency of ≥97% identity, in both strand orientations, with maxaccepts=1 and maxrejects=0.

### Data Availability

Full-length Sanger Sequenced 16S rRNA gene sequences are available on NCBI GenBank, accession numbers: MK875836 - MK875967. Illumina data from the 55 cm Oracle Ridge community are available on the NCBI SRA, accession numbers: SRR9172130-SRR9172198.

## Acknowledgements

The authors would like to thank Nathan Abramson, Jasper Bloodsworth, Brenna Bourque, Amanda Howe, Bridget Taylor, and H. James Tripp for assisting with sample collection, culture maintenance, and DNA extractions relevant to this work. Funding from this work came from startup funds provided to PC from the University of Arizona’s Technology and Research Initiative Fund (the Water, Environmental, and Energy Solutions initiative), and seed grants from the Center for Environmentally Sustainable Mining and The University of Arizona College of Agriculture and Life Sciences. Work at JCVI was supported by P01AI118687.

## Supplementary Table and Figure Captions

**Supplementary Table 1.** Artificial Subterranean Growth Medium (ASM) -low and -high formulations including concentrations of carbon sources, amino acids, vitamins, trace metals, amino acids, and inorganic nutrients.

**Supplementary Figure 1: The soils used for cultivation were oligotrophic.** (A) points are the measured soil organic carbon content of 184 soil samples collected from shallow subsurface pits located across the United States (from ref. (11)). (B) points are the measured NO^3^-N content of 121 soils (113 surface soils) collected across the United States (from refs (91, 92)). The organic carbon percent and NO^3^-N measured in our 55 cm sample from Oracle Ridge is illustrated with a red dashed line in both panels. Our sample falls on the low end of both organic carbon content and NO^3^-N content across this broad selection of soils.

**Supplementary Figure 2: Alphaproteobacterial IQ-TREE Maximum Likelihood Full Length 16S rRNA gene sequence phylogeny.** The tree was constructed using a GTR+F+I+G4 nucleotide substitution model iterated over 10,000 ultrafast bootstraps. The scale bar indicates the FigTree proportionally transformed branch length. The 16S rRNA gene sequence from *E. coli* K-12 was used as an outgroup. Clade bootstrap support is indicated by colored nodes as indicated in the legend. This tree includes 16S rRNA gene sequences that amplified from all pure alphaproteobacterial cultures isolated in this study (bolded black leaves denoted with ‘AZCC’), as well as their best sequence matches to NCBI microbial genomes (dark blue leaves), NCBI 16S rRNA gene sequences from cultured isolates (orange leaves), and NCBI 16S rRNA gene sequences from environmental clones (green leaves). The growth medium concentration on which each AZCC culture was isolated are indicated by colored bars on the right side of the phylogenetic tree (red for ASM-high and blue for ASM-low).

**Supplementary Figure 3: Actinobacterial IQ-TREE Maximum Likelihood Full Length 16S rRNA gene sequence phylogeny.** The tree was constructed using a SYM+R10 nucleotide substitution model iterated over 10,000 ultrafast bootstraps. The scale bar indicates the FigTree proportionally transformed branch length. The 16S rRNA gene sequence from *B. subtilis* DSM-10 was used as an outgroup. Clade bootstrap support is indicated by colored nodes as indicated in the legend. This tree includes 16S rRNA gene sequences that amplified from all pure actinobacterial cultures isolated in this study (bolded black leaves denoted with ‘AZCC’), as well as their best sequence matches to NCBI microbial genomes (dark blue leaves), NCBI 16S rRNA gene sequences from cultured isolates (orange leaves), and NCBI 16S rRNA gene sequences from environmental clones (green leaves). The growth medium concentration on which each AZCC culture was isolated are indicated by colored bars on the right side of the phylogenetic tree (red for ASM-high and blue for ASM-low). Actinobacterial classes are indicated by colored vertical bars to the right of the tree figure: Thermoleophilia (purple), Acidimicrobiia (teal), and Actinobacteria (Orange-Red).

**Supplementary Figure 4: Distribution of nucleic acid fluorescence values by Phylum (a) and Genus (b).** Data are natural logarithm-transformed nucleic acid fluorescence values of fixed and SYBR Green I-stained stationary phase cultures. Plots show all genera with n ≥ 3 cultures in one or both culture media types. Shaded regions illustrate the distribution of fluorescence values within culture media groups.

